# VPS41 deletion triggers progressive loss of insulin stores and downregulation of beta-cell identity

**DOI:** 10.1101/2024.04.17.589848

**Authors:** Belinda Yau, Yousun An, Mark Germanos, Patricia Schwarzkopf, A Gabrielle van der Kraan, Mark Larance, Hayley Webster, Christian Burns, Cedric S Asensio, Melkam A Kebede

**Author notes:** These authors contributed equally (Belinda Yau, Yousun An).

## Abstract

Vacuolar protein sorting-associated protein 41 (VPS41) has been established as a requirement for normal insulin secretory function in pancreatic beta-cells. Genetic deletion of VPS41 in mouse pancreatic beta-cells results in diabetes, though the mechanisms are not understood. Presently, we show that VPS41 deletion results in rapid mature insulin degradation and downregulation of beta-cell identity. This phenotype is observed *in vivo,* with VPS41KO mice displaying progressive loss of insulin content and beta-cell function with age. In acute VPS41 depletion *in vitro*, the loss of insulin is associated with increased degradative pathway activity, increased Adapter Protein 3 complex colocalisation with lysosomes, increased nuclear localisation of transcription factor E3, and downregulation of PDX1 and INS mRNA expression. Inhibition of lysosomal degradation rescues the rapidly depleted insulin content. These data evidence a VPS41-dependent mechanism for both insulin content degradation and loss of beta-cell identity in beta-cells.

## Introduction

Regulation of insulin content in pancreatic beta-cells is at the core of beta-cell function. It relies on a carefully curated balance between insulin secretory granule (ISG) biosynthesis, storage, degradation, and secretion[1]. In conditions of glucose abundance when more insulin secretion is required, insulin degradation is decreased[2] and insulin synthesis is increased [3]. This is characterised in the beta-cell by enhanced insulin promoter activation, elevated INS1 gene expression [3], and expansion of the endoplasmic reticulum and Golgi apparatus to accommodate increased ISG formation[4]. Intracellular ISG storage is also higher, reflected in increased beta-cell mass[4]. On the contrary, in conditions of reduced secretory need, ISG synthesis is downregulated, and insulin storage is minimized. Insulin granule degradation to control insulin content is mediated through multiple cellular pathways; crinophagy – direct insulin granule to lysosome fusion to degrade granule contents[5], conventional macroautophagy – in which insulin granules are contained within autophagosomal bodies for lysosomal fusion – or microautophagy[6] – in which insulin granules are autophagocytosed by the lysosomal compartment, also appear to contribute to insulin granule turnover[7]. There also exist specialized pathways to degrade ISGs known as stress/starvation-induced nascent granule degradation (SINGD), which are activated to preserve cellular resources[8, 9].

Vacuolar protein sorting-associated protein 41 (VPS41) is a core component of the homotypic fusion and vacuole protein sorting (HOPS) tethering complex, which functions as a mediator of vesicle fusion between intracellular degradative compartments of the cell[10–12]. Independent of the HOPS complex, VPS41 has additional roles in protein trafficking between vacuolar compartments such as the lysosome and endosome through interactions with Rab7[13], Adapter Protein 3 (AP3)[14, 15] and Rab5[16], though some of these pathways have yet to be verified in mammalian cells. More recently, a role for VPS41 in the regulated secretory pathway has been identified, implicating the protein in granule biogenesis for the first time[14].

Human VPS41 mutations have recently been characterised[17], with these extending to clinically relevant single nucleotide polymorphisms (SNPs) in VPS41 causing neurodegenerative pathologies in human families[18]. Inherited biallelic loss of function VPS41 variants have also been evidenced to cause progressive dystonia and cerebellar dysfunction in patients, with implications for impaired lysosomal function[18, 19]. Importantly, these neuropathologies appear to be driven by dysfunction of the HOPS complex, in which a VPS41 mutation results in a HOPS loss of function phenotype.

In pancreatic beta-cells, VPS41 deletion and SNPs exhibited similar phenotypes to those described in the brain[17, 20]. Dissolution of HOPS complex function by either VPS41 or VPS39 – another core component of HOPS – depletion resulted in impaired endolysosomal fusion, and consequently defective cargo degradation[20]. Importantly, our initial study in insulinoma INS1 beta-cells established the existence of a separate pool of HOPS-independent VPS41 protein that also contributes to ISG biogenesis, and this VPS41 pool remained stable despite HOPS destabilization when VPS39 alone was depleted. We showed significant changes to insulin granule morphology and composition occured in CRISPR-Cas9-generated VPS41 knockout beta-cells[20], building on prior findings that VPS41 may exert a role as a coat protein for granule formation[14]. Most strikingly, a S284P single nucleotide polymorphism in VPS41 was sufficient to disrupt HOPS while keeping VPS41’s function in the insulin secretory pathway intact, suggesting that VPS41 has both HOPS-dependent and - independent functions in the beta-cell. Importantly, we also determined that *in vivo* genetic deletion of VPS41 specifically in beta-cells using the *Ins1-Cre* knock-in mouse line[21] resulted in an insulin-sensitive diabetic phenotype in mice, characterised by insulin insufficiency and extreme hyperglycaemia[20]. These findings were a crucial step in defining the relationship between VPS41 and the insulin secretory pathway.

In our present study we attempt to disentangle the acute and chronic effects of VPS41 deletion in pancreatic beta-cells using an siRNA-mediated knockdown of VPS41 in insulinoma cells (VPS41KD), and the characterisation of younger VPS41KO mice. We show that beta-cell specific deletion of VPS41 in mice results in a worsening reduction of insulin content and glucose tolerance with age. In INS1 beta-cells, VPS41 depletion by silencing RNA also significantly reduces insulin content by 48 h, and this can be rescued by pre-treatment with lysosomal inhibitors. In VPS41KO islets, PDX1 downregulation precedes insulin loss, which results in beta-cell dedifferentiation in the chronic state. Altogether these data demonstrate a requirement for VPS41 in insulin content maintenance in beta-cells.

## Experimental Procedures/Methods/Materials and Methods

### Cells

INS-1 cells (gifted from Dr. Peter Arvan, University of Michigan, Ann Arbor, MI) were maintained in RPMI (with 10% FBS, 1 mmol/L sodium pyruvate, 50 µmol beta-mercaptoethanol, 1% penicillin-streptomycin) at 5% CO_2_ and 37°C. VPS41KO cells were generated as previously described[20].

### Plasmids

VPS41 siRNA for knockdown experiments was commercially generated (Thermo Fisher Silencer Select siRNAs) using siRNA Assay ID s157874, alongside Silencer Select Negative Control No.1 (4390843). proinsulin-msGFP2 was generated from pQE-60NA-proinsulin-msGFP2 (Addgene, #160466) and verified by Sanger sequencing (Garvan Molecular Genetics, Australia). proCpepSNAP was a gift from Samuel Stephens[22].

### Degradation Inhibition

Chloroquine (C6628), MG132 (M8699), E64D (E3132) and ammonium chloride (326372) were all obtained from Sigma-Aldrich.

### qPCR

Cell samples were frozen in 500 µL of RLT buffer containing 1 % β-mercaptoethanol. RNA extraction was performed using a RNeasy Mini Kit (Qiagen) as per manufacturer’s protocol, and cDNA was obtained using 1 µg RNA and the one-step iScript Reverse Transcription Supermix (Biorad). qPCR was performed using SYBR SELECT Master Mix on a Roche Lightcycler 480 II and relative expression was calculated using the ΔΔCt method using GAPDH as a housekeeping gene. Primers used were *Pdx1*: forward 5’AAATCCACCAAAGCTCACGC-3’, reverse, 5’AAGTTGAGCATCACTGCCAGC-3’, *Gapdh*: forward, 5’-CAAAATGGTGAAGGTCGGTGTG-3’, reverse,5’-TGATGTTAGTGGGGTCTCGCTC-3’, *Ins2*: forward, 5’-GCAGGTGACCTTCAGACCTT-3’, reverse, 5’-CAGAGGGGTGGACAGGGTAG-3’. *Mafa*: forward, 5’-AGGAGGAGGTCATCCGACTG-3’, reverse, 5’-CTTCTCGCTCTCCAGAATGTG-3’. *Bhlhe40*: forward, 5’-CCGTGTGGTCTCTGAACTCC-3’, reverse, 5’-CACCCTTCTCCAATTCGCCT-3’.

### Mass spectrometry-based proteomics

Protein samples for bottom-up proteomics analysis were prepared as describe previously[23]. Briefly, proteins were extracted in a buffer containing 4% sodium deoxycholate (SDC), 40 mM chloroacetamide, 10 mM TCEP and 0.1 M Tris-HCl (pH 8.0) in water. Samples were then incubated at 95°C for 10 min at 1000 rpm in a Thermomixer-C (Eppendorf). Samples were diluted to a final concentration of 1% SDC with water and 10 ug of protein digested overnight with 200 ng of MS-grade trypsin (Sigma) in 50 mM acetic acid at 37°C, on a Thermomixer-C (Eppendorf) at 1000 rpm. Samples were mixed with ethyl acetate (50% final concentration, v/v), vortexed until SDC precipitates dissolved. SDB-RPS StageTips were prepared, sample purification performed as described previously[23]. Peptides were reconstituted with 5 % formic acid in MS-grade H2O, sealed and stored at 4 °C until LC-MS/MS acquisition.

Using a ThermoFisher RSLCnano ultrahigh performance liquid chromatograph, peptides in 5 % (v/v) formic acid (3 µL injection volume) were injected onto a 50 cm x 75 µm C18Aq (Dr Maisch; 1.9 µm) fused analytical column with a ∼10 µm pulled tip, coupled online to a nanospray electrospray ionization source. Peptides were resolved over gradient from 5 – 40 % acetonitrile for 70 min, with a flow rate of 300 nL min-1 (Capillary flow). Electrospray ionization was done at 2.3 kV. Tandem mass spectrometry analysis was carried out on a Exploris or Lumos Tribrid mass spectrometer (ThermoFisher) using data-independent acquisition (DIA). DIA was performed as previously described using variable isolation widths for different m/z ranges 114. Stepped normalized collision energy of 25 ± 10 % was used for all DIA spectral acquisitions.

Raw MS data were analyzed using quantitative DIA proteomics software DIA-NN (v. 1.8). Complete mouse proteome databased from UNIPROT was used for neural network generation, with deep spectral prediction enabled. Protease digestion was set to trypsin (fully specific) allowing for two missed cleavages and one variable modification. Oxidation of Met and acetylation of the protein N-terminus were set as variable modifications. Carbamidomethyl on Cys was set as a fixed modification. Match between runs and remove likely interferences were enabled. Neural network classifier was set to double-pass mode. Protein interferences were based on genes. Quantification strategy was set to any liquid chromatography (LC, high accuracy). Cross-run normalization was set to RT-dependent. Library profiling was set to smart profiling. The mass spectrometry proteomics data have been deposited to the ProteomeXchange Consortium via the EBI PRIDE partner repository with the dataset identifier PXD055718, Username, Password: gD3uJD1ln3Ry.

### Mice

Ins1-Cre control (CRE)[21], and beta-cell specific VPS41 heterozygous (βVPS41HET) and homozygous (βVPS41KO) mice were bred at Australian BioResources (Garvan institute of Medical Research) and transferred to the Charles Perkins Centre (The University of Sydney) for experimental procedures. Mice were fed a standard laboratory chow (13% calories from fat, 65% carbohydrate, and 22% protein) from Gordon’s Specialty Stock Feeds (Yanderra, New South Wales, Australia). VPS41*fl/fl* mice (26) were crossed with Ins1-Cre mice (strain 026801; The Jackson Laboratory) to generate third generation VPS41*fl/fl*, VPS41*fl/+* and *Ins1-cre*/+ mice and their littermate controls for experiments. Male and female mice were used where indicated. Mice were maintained on a 12-h light/dark cycle (0700/1900 h) with *ad libitum* access to food and water. Experiments were carried out in accordance with the National Health and Medical Research Council (Australia) guidelines for animal research and approved by The University of Sydney Animal Experimentation Ethics Committees (Camperdown, New South Wales, Australia).

### Mouse metabolic phenotyping

Magnetic resonance imaging (MRI) was conducted (EchoMRI 900) to determine lean and fat mass and mice were administered glucose (2 g / kg lean mass) by oral gavage. Mice were fasted for 5 hours prior to oral-GTT. Blood glucose was measured immediately prior to and 15, 30, 45, 60 and 90 min after glucose injection with ACCU-CHEK Performa II Strips (Roche, Basel, CH). Blood was collected immediately prior to and 15 minutes after glucose injection for plasma insulin measurements.

### Islet Isolation and Glucose-induced insulin secretion assays

Islets were harvested from mice as previously described[24]. Glucose-induced insulin secretion assays were performed in isolated islets as previously described[24]; islets were placed into 2.8 mM glucose Krebs Ringer buffer supplemented with 10 mM HEPES (KRBH) for 1 hour prior to assay as pre-basal. New KRBH media containing 2.8 mM was added, then collected after 1 hour (basal secretion). Islets were then stimulated with KRBH containing 16.7 mM glucose and supernatant collected after 1 hour (stimulated secretion). Islets were then washed in PBS and lysed by sonication in Islet Lysis Buffer (100mM Tris pH7.4, 300mM NaCl, 10mM NaF) to measure total islet insulin content.

### Insulin measurements

Plasma insulin measurements were obtained using an Ultra-Sensitive Insulin ELISA Assay (Crystal Chem). *Ex vivo* islet and INS1 cell glucose-stimulated insulin secretion assay insulin measurements were obtained using a rapid homogenous time-resolved fluorescence (HTRF) Insulin Assay kit (Cisbio Assay).

### Immunofluorescent staining

Paraffin-embedded pancreas tissue was processed as previously described [20]. Immunofluorescent staining for proinsulin, insulin, glucagon, TFE3, and DAPI was performed as previously described[25]. Pancreas slices and cell coverslips were washed and mounted with ProLong Diamond Antifade containing DAPI (Invitrogen). Immunofluorescently-stained images from both pancreas tissue and cells were obtained using a white-light laser equipped Leica SP8 confocal microscope at 40X and 63X magnification respectively, using Leica LAS X software.

### SDS-PAGE

Proteins in whole-cell lysates and fractions were separated using MES buffer SDS-PAGE under denaturing conditions on 4 - 12% Bis-Tris Bolt^TM^ polyacrylamide gels (Invitrogen), then transferred to methanol-activated PVDF membranes. The modified Western blotting method recently described by Okita et al.[26] was used to immunoblot for proinsulin and insulin. Control proteins (beta-actin and GAPDH) were probed on the same blots as target proteins using alternate species antibodies.

### Antibodies

The following commercial antibodies and their sources are listed: VPS41 (sc-377118, SCBT), LC3A/B (12741, CST), LAMP1 (ab24170, Abcam), Insulin (IR00261-2, Agilent DAKO), Proinsulin (GS-9A8, DSHB), Chromogranin A (NB120-15160SS, Novus Biologicals), Rab5 (3547, CST), Beta-actin (A5441, Sigma), phospho-eIF2a (3398, CST), total-eIF2a (5324, CST), phospho-IRE1-alpha (PA1-16927, Invitrogen), total IRE1-alpha (3298, CST), Beclin-1 (3495, CST), Atg12 (4180, CST), CHOP (2895, CST), BiP/GRP78 (3177, CST), phospho-JNK (4668, CST), total-JNK (67096, CST), mTORC1 (2983, CST), TFE3 (MA537818, Invitrogen).

### Analysis and Statistics

Prior to analysis, immunofluorescence images for colocalisation analysis were deconvoluted using Huygens Professional software (Scientific Volume Imaging). Immunofluorescence image analysis for fluorescence intensity, colocalisation and western blot quantification was performed in FIJI (Image J). Proteomic analysis was performed in Perseus (v1.6.15 MaxQuant). Statistical analyses were performed in Prism (GraphPad Prism 8).

## Results

### *In vivo* VPS41 deletion in beta-cells causes progressive degranulation with age

Previously, mixed-sex beta-cell specific VPS41 KO mice (βVPS41KO) were characterised at 8 – 9 weeks and 15 - 16 weeks age[20], demonstrating a severe loss of insulin content and insulin secretory capacity, resulting in significantly elevated blood glucose levels with no changes in insulin sensitivity. When comparing Cre controls to beta-cell specific VPS41 heterozygous mice (βVPS41HET) at 12 weeks, no differences were observed in fasting blood glucose (Supplemental Figure 1A), glucose tolerance (Supplementary Figure 1B), or VPS41 protein levels (Supplemental Figure 1C and D). As a result, βVPS41HET mice were used as controls in subsequent experiments.

Characterisation of 5 - 6 week-old control and βVPS41KO female mice revealed no differences in body composition (Figure 1A), but βVPS41KO mice exhibited significantly elevated fasting blood glucose levels (Figure 1B). Unlike in older βVPS41KO mice[20], this increase was not accompanied by a significant reduction in fasting insulin levels (Figure 1C). Blood glucose levels and the corresponding area under the curve during an oral glucose tolerance test (oGTT) were significantly higher in βVPS41KO mice (Figure 1D and E), though insulin levels during the oGTT were not significantly different (Figure 1F). When the incremental area under the curve for the oGTT was calculated (Figure 1G and H), no differences were observed between control and βVPS41KO mice, suggesting the elevated fasting glucose was the primary factor contributing to the impairment in glucose tolerance. In *ex vivo* analysis of isolated islets, both control and βVPS41KO islets showed significant response to 16.7 mM (high) glucose stimulation (Figure 1I), but βVPS41KO islets secreted significantly less insulin compared to controls when normalised to DNA content (Figure 1J). No differences in insulin content were detected in islets isolated from female mice. (Figure 1K and L).

**Figure 1.**
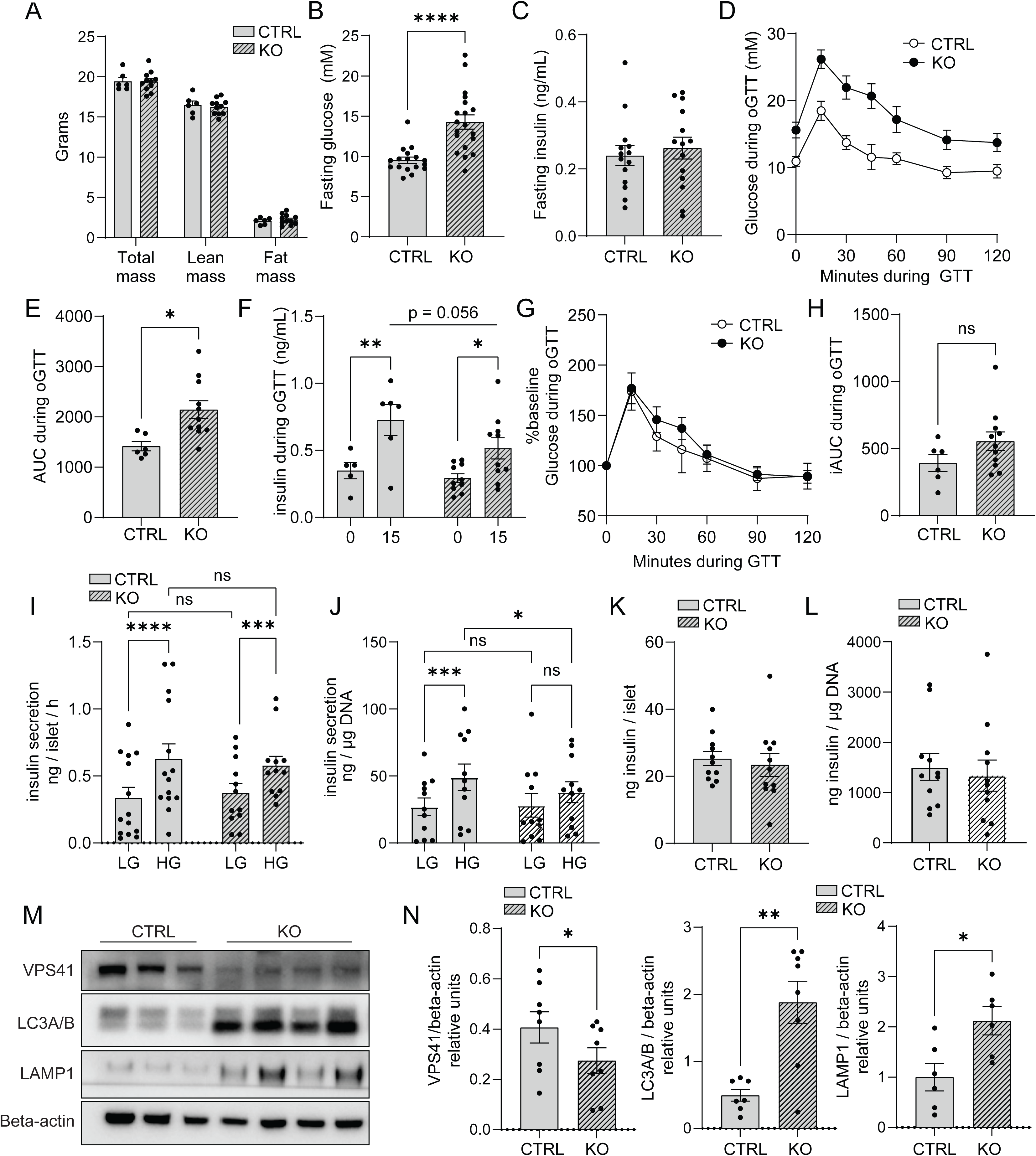
Female βVPS41KO mice exhibit a mild insulin-deficiency phenotype with increased islet expression of degradation-associated proteins. Total, lean and fat mass **(A)**, fasting blood glucose **(B)** and fasting blood insulins **(C)** in female βVPS41KO (KO) and βVPS41HET (CTRL) mice at 5 - 6 weeks of age. Glucose measurements **(D)**, area under the curve **(E)**, and insulin measurements from female CTRL and KO mice during oral glucose tolerance testing **(F)**. Glucose measurements expressed as a percent of fasting glucose **(G)** and incremental area under the curve **(H)** of oral glucose tolerance test curves from female KO and CTRL mice. *Ex vivo* glucose-stimulated insulin secretion assay to measure insulin secretion at low (2.8mM) and high (16.7mM) glucose **(I)** and total insulin content per islet **(K)** and insulin secretion **(J)** and islet content **(L)** expressed relative to DNA content in female KO and CTRL mice. Representative western blot **(M)** and quantitative expression **(N)** of VPS41, LC3A/B, and LAMP1 relative to beta-actin in *ex vivo* pancreatic islets from female KO and CTRL mice at 6 - 7 weeks age. Data expressed as mean ± SEM, *p < 0.05, **p < 0.01 by unpaired t-test.

Phenotypic analysis of male control and βVPS41KO mice showed results consistent with previous observations in older βVPS41KO mice. There were no change in body composition (Supplemental Figure 2A), but fasting glucose levels were markedly elevated (Supplemental Figure 2B), along with glucose intolerance observed through oGTT (Supplemental Figure 2C and D). Both the area under the curve and the incremental area under the curve were also significantly increased (Supplemental Figure 2E and F), which corresponded to severely impaired insulin secretion during oGTT (Supplemental Figure 2G). Additionally, *ex vivo* islets from male βVPS41KO mice exhibited significantly reduced insulin secretion in response to glucose stimulation (Supplemental Figure 2H and I), as well as a marked decrease in islet insulin content (Supplemental Figure 2J and K).

#### *In vivo* VPS41 deletion upregulates degradative pathways in islets

Western blotting analysis of *ex vivo* islets from 6-week-old male and female βVPS41KO mice revealed significant reduction in VPS41 protein levels, along with a marked increase in LAMP1 and LC3A/B expression - proteins associated with lysosomes and autophagosomes, respectively (Figure 1M and N). Similar results were observed in male βVPS41KO mice (Supplemental Figure 3A and B), where CHOP expression was elevated and phospho-JNK expression was increased exclusively in male mice (Supplemental Figure 3A and C). No significant differences were found in other proteins related to endoplasmic stress, such as Beclin-1, phospho-IRE-1, phospho-eIF1-alpha, Atg12, and BiP (Supplementary Figure 3A and C). Interestingly, ovariectomy in 4 – 5-week-old female control and βVPS41KO mice did not affect blood glucose levels (Supplemental Figure 3D), nor did female ovariectomized mice exhibit blood glucose levels similar to older male mice (Supplemental Figure 2B), indicating that the sexually dimorphic response is not solely driven by estrogen.

We then shifted our focus to female βVPS41KO mice, using them as a model to study the early stages of VPS41-induced insulin deficiency, before an overt loss of insulin content. Fed blood glucose concentrations were compared between 6-week and 16-week-old control and βVPS41KO mice, revealing an age-dependent dysregulation of glucose homeostasis in βVPS41KO mice only (Figure 2A). Next, we performed proteomic analysis of islets from 6 week and 16-week female control and βVPS41KO mice and showed age-dependent changes in the islet proteome of βVPS41KO mice. In 6-week-old βVPS41KO mice, 11% of the islet proteome (761 proteins) was differentially expressed compared to controls, while in 16-week-old βVPS41KO mice, this increased to 43% (3084 proteins) (Figure 2B). Gene Ontology (GO) pathway analysis of the differentially expressed pathways revealed that upregulated pathways related to degradation, such as *macroautophagy*, *lytic vacuole organisation* and *lysosomal transport*, were already present at 6 weeks and persisted through 16 weeks (Figure 2C).

**Figure 2.**
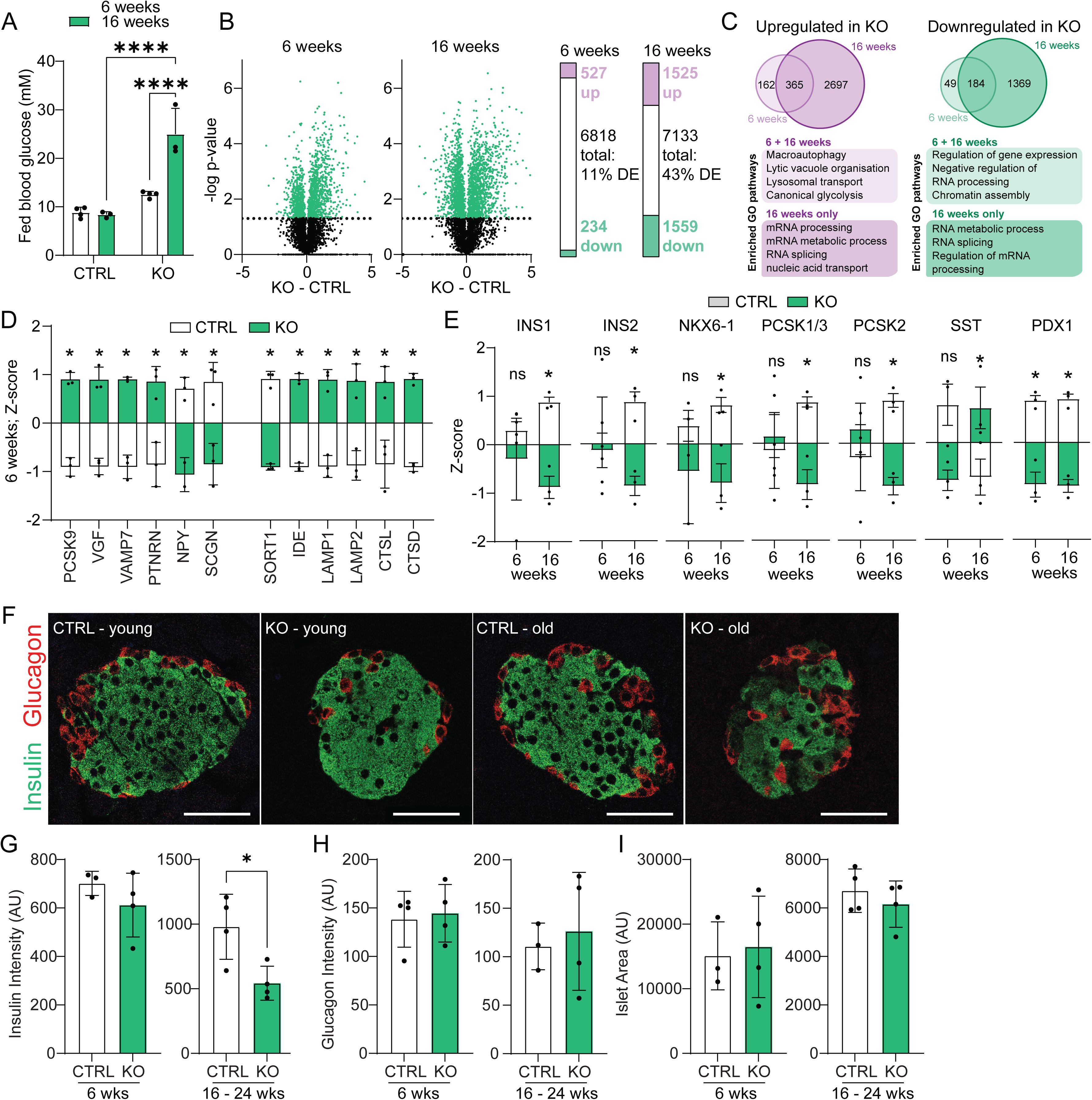
Islet proteomics of VPS41 beta-cell specific KO female mice at 6 and 16 weeks of age. Blood glucose levels of female βVPS41HET (CTRL) and βVPS41KO (KO) mice at 6 and 16 weeks of age **(A)**. Proteomic analysis (n = 3) of CTRL and KO female mice at 6 and 16 weeks of age identified 6818 and 7133 proteins respectively, with differentially expressed proteins representing 11% and 43% of the total respective proteome **(B)**. Gene Ontology pathway enrichment was performed on upregulated and downregulated protein sets and compared between 6- and 16-week-old mice **(C)**. Protein expression levels of key insulin granule, protein degradation and trafficking-associated genes were extracted from the proteomics data to highlight the CTRL/KO comparisons at 6 weeks **(D)**. Comparison of protein expression of INS1, INS2, PDX1, NKX6.1, NKX2.2, PCSK1/3, PCSK2 and SST in 6- and 16-week-old female mice **(E)**. Immunofluorescent staining of insulin and glucagon in paraffin-embedded pancreatic islet slices of female CTRL and KO mice at 6 – 7 weeks, or 16 – 24 weeks of age **(F)**. Quantification of insulin fluorescence intensity **(G)**, glucagon fluorescence intensity **(H)** and pancreatic islet area **(I)**. 3 – 4 islets per mouse analysed, each dot represents an average mean intensity or area of one mouse. Bars represent mean ± SEM. *p < 0.05, ****p < 0.0001 by unpaired t-test or one-way ANOVA.

Interestingly, pathways that were downregulated at 6 weeks and persisted at 16 weeks include the *regulation of gene expression, negative regulation of RNA processing* and *chromatin assembly* (Figure 2D). In the islets of 6-week-old mice, we observed changes in the expression of dysregulated granule-associated proteins, such as increased expression of receptor-type tyrosine-protein phosphatase-like N (PTPRN) and VGF protein (VGF), alongside decreased levels of neuropeptide Y (NPY) and secretogranin (SCGN). Additionally, there was an upregulation of autophagy-associated vesicle-associated membrane protein 7 (VAMP7), lysosomal proteins (LAMP1, LAMP2), insulin degrading enzyme 1 (IDE1) and cathepsins (CTSL, CTSD, Figure 1D). Finally, while insulin (INS1, INS2) levels were not significantly altered at 6 weeks, they were significantly downregulated by 16 weeks. This reduction was preceded by a notable decrease in the transcription factor PDX1 at 6 weeks (Figure 2E). Further significant downregulation of transcription factor NKX6.1, prohormone convertases 1 (PCSK1/3) and 2 (PCSK2) along with upregulation of somatostatin (SST) at 16 weeks, suggesting potential alterations to islet cell function with age. All differentially expressed proteins in both 6- and 16-weeks KO and CTRL mice are summarized in Supplemental Table 1.

Pancreas sections from female CTRL and βVPS41KO mice, aged 6 – 7 weeks and 16 – 24 weeks, were stained for insulin and glucagon. We observed a significant reduction in mean insulin intensity in βVPS41KO mice compared to CTRL, but this difference was only apparent in the older mice (Figure 2F and 2G). In contrast, there was no significant difference in mean glucagon intensity between young and old mice (Figure 2F and 2H), regardless of genotype. Additionally, mean islet area did not differ between βVPS41KO and CTRL mice at either age (Figure 2I). These findings suggest a specific loss of insulin in the pancreatic islets of older βVPS41KO mice.

### Acute depletion of VPS41 in INS1 beta-cells results in loss of insulin stores

To further explore the effects of acute and chronic VPS41 deletion in beta-cells, we established a VPS41 depletion model using siRNA-mediated VPS41 knockdown in WT INS1 cells (VPS41KD). Over a 72-hour period, VPS41 depletion led to a corresponding depletion of insulin granules, as indicated by decreases in insulin and chromogranin A (CgA) levels, without affecting lysosomes (LAMP1) or endosomes (Rab5, Figure 3A) at these time points. At 24 hours post-transfection, VPS41 levels were reduced to 33% of the non-targeting transfection controls (Supplemental Figure 4A), and both proinsulin and insulin levels were significantly lower (Supplemental Figure 4B and C). By 48 hours post-transfection, VPS41 levels dropped to 22 % of controls (Figure 3B) and insulin levels were 18 % of controls as measured by SDS-PAGE immunoblotting (Figure 3C). Proinsulin levels also decreased to 61% of controls at the 48 hour (Figure 3D) though this reduction was less pronounced than for insulin, resulting in a significant increase in the proinsulin/insulin ratio (Figure 3E). This suggests a disruption in the balance between mature insulin granules and immature insulin granules, and/or proinsulin synthesis. This was further supported by immunofluorescence analysis, which showed that proinsulin and insulin levels were 47 % of control (Supplementary Figure 4D and E).

**Figure 3.**
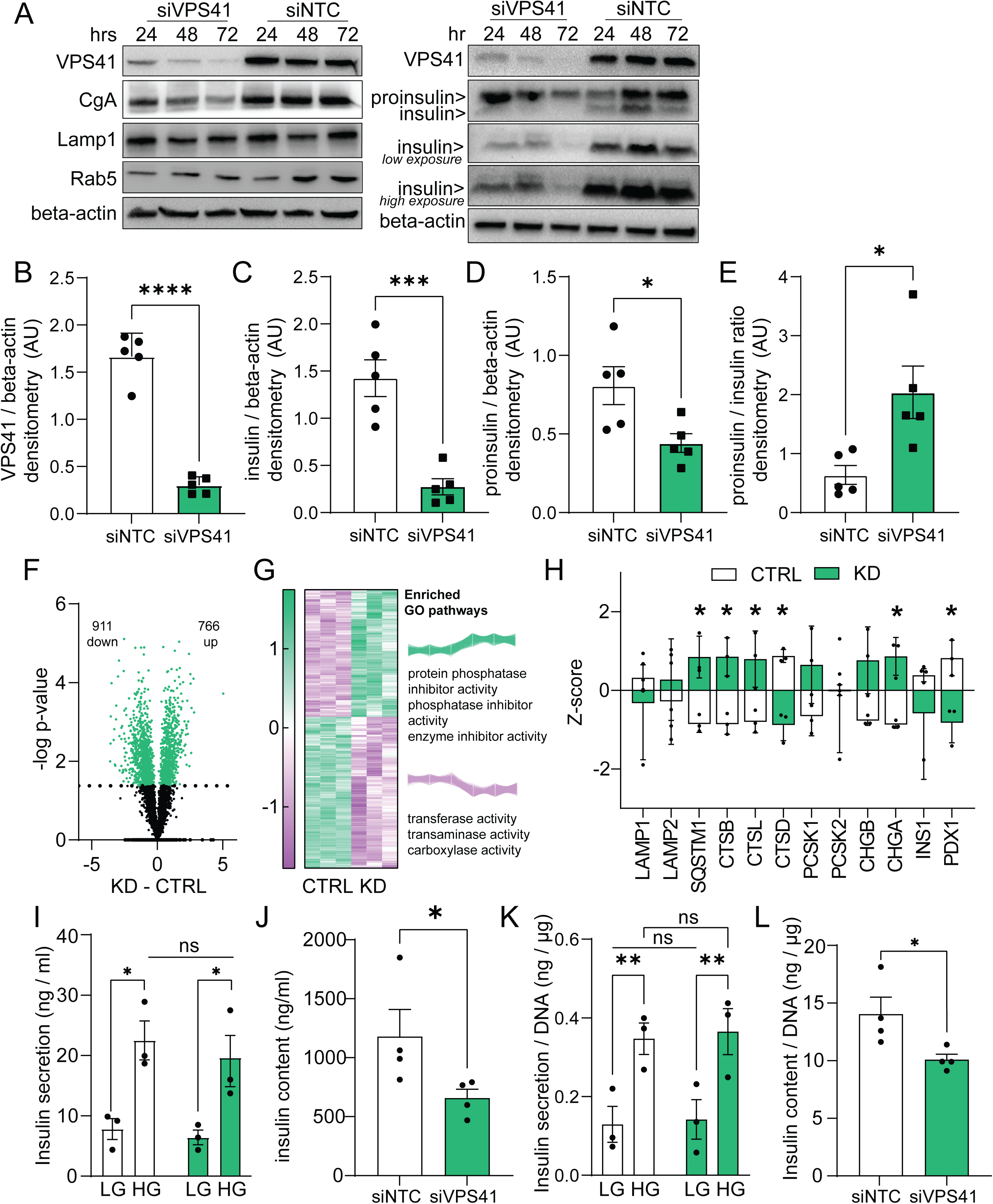
Acute VPS41 depletion in INS1 cells by siRNA-mediated knockdown results in diminished insulin content. Protein expression of VPS41, chromogranin A, Lamp1, Rab5, insulin, proinsulin, and beta-actin after siRNA-mediated knockdown of VPS41 (50 pmol) and non-targeting control (NTC) siRNA over a 72-hour time course **(A)**. VPS41 **(B)**, insulin **(C)**, proinsulin **(D)**, and proinsulin over insulin ratio **(E)** protein expression relative to beta-actin loading control at 48 hours post-siRNA-mediated KD of VPS41 (50pmol) in INS1 cells. Proteomics analysis (n = 3) of VPS41KD cells at 72 h post-transfection identified 1677 differentially expressed proteins **(F)**, from which Gene Ontology pathway enriched was performed **(G)**. Expression levels of key degradation-associated and insulin secretory pathway-associated proteins were extracted from the proteomics data to highlight these comparisons **(H)**. Glucose-stimulated insulin secretion assay to measure insulin secretion at low (2.8mM) and high (16.7mM) glucose **(I)** and total insulin content **(J)** normalised to DNA **(K** and **L)** in VPS41KD and NTC siRNA control cells at 72 hours post-transfection. Data expressed as mean ± SEM, *p < 0.05, **p < 0.01, ***p < 0.001, ****p < 0.0001 by unpaired t-test or two-way ANOVA.

When we then performed proteomic analysis of VPS41KD cells we identified 1677 proteins differentially expressed compared to non-targeting siRNA controls (Figure 3F). GO pathway analysis revealed upregulation in pathways related to *protein phosphatase inhibitor activity* and *enzyme inhibitor activity* (Figure 3G). While the expression of LAMP1 and LAMP2 remained unchanged, we observed significant upregulation of the autophagosome cargo marker sequestosome 1 (SQSTM1) and cathepsins L and B (CSTL, CSTB) in VPS41KD cells. Conversely, cathepsin D (CSTD) was decreased. Similar to 6-week-old female βVPS41KO islets, there was no significant change in (pro)insulin (INS1), but PDX1 levels were significantly reduced (Figure 3H). Surprisingly, despite no overt changes in insulin granule-associated proteins such as prohormone convertases (PCSK1, PCSK2) or chromogranin B (CHGA), we observed an increase in Chromogranin B (CHGB) (Figure 3H). Additionally, expression of other core HOPS subunits (VPS11, VPS16, VPS18, VPS33a) was not detected by mass spectrometry in VPS41KD cells, suggesting their levels might also have been significantly reduced due to destabilization of the HOPS complex (Supplemental Figure 4F). A complete list of differentially expressed proteins in VPS41KD is provided in Supplemental Table 2. Finally, a glucose-stimulated insulin secretion assay performed on VPS41KD cells 72 hours post-transfection revealed no defect in insulin secretion when cells were stimulated with 16.7 mM glucose (Figure 3I and K). However, there was a significant reduction in insulin content (Figure 3J and L), indicating that acute VPS41 deletion primarily affects insulin stores rather than the secretory function.

### Insulin granules are degraded through lysosomal pathways in VPS41-depleted INS1 beta-cells

Insulin granules typically have a half-life of 3 to 5 days within beta-cells. Thus, the observed depletion of insulin stores in VPS41-depleted cells within just 24 to 48 hours suggest that insulin granules are being redirected towards degradative pathways. This hypothesis is supported by the increased expression of degradation pathway-associated proteins observed in both βVPS41KO islets (Figure 2D) and VPS41KD cells (Figure 3H).

To further investigate this hypothesis, we pre-treated INS1 cells for 18 hours with various degradation inhibitors, including lysosomal inhibitors chloroquine and ammonium chloride, cysteine protease inhibitor E64D, and proteosome inhibitor MG132, before performing VPS41 siRNA-mediated knockdown (Figure 4A and B). Our results showed that pretreatment with chloroquine, E64D and ammonium chloride was able to restore insulin levels to those of control cells, as measured by both Western blot (Figure 4A and C) and ELISA (Supplementary Figure 4G). This suggest that the loss of insulin is occurring through lysosomal degradation pathways. In contrast, pretreatment with MG132 did not have this effect, indicating that pro/insulin loss is not occurring through proteasomal degradation. Additionally, we investigated whether insulin was redirected to constitutive secretion within the first 24 hours of siRNA treatment, but we did not observe increased insulin secretion in the media of VPS41KD cells (Figure 4D).

**Figure 4.**
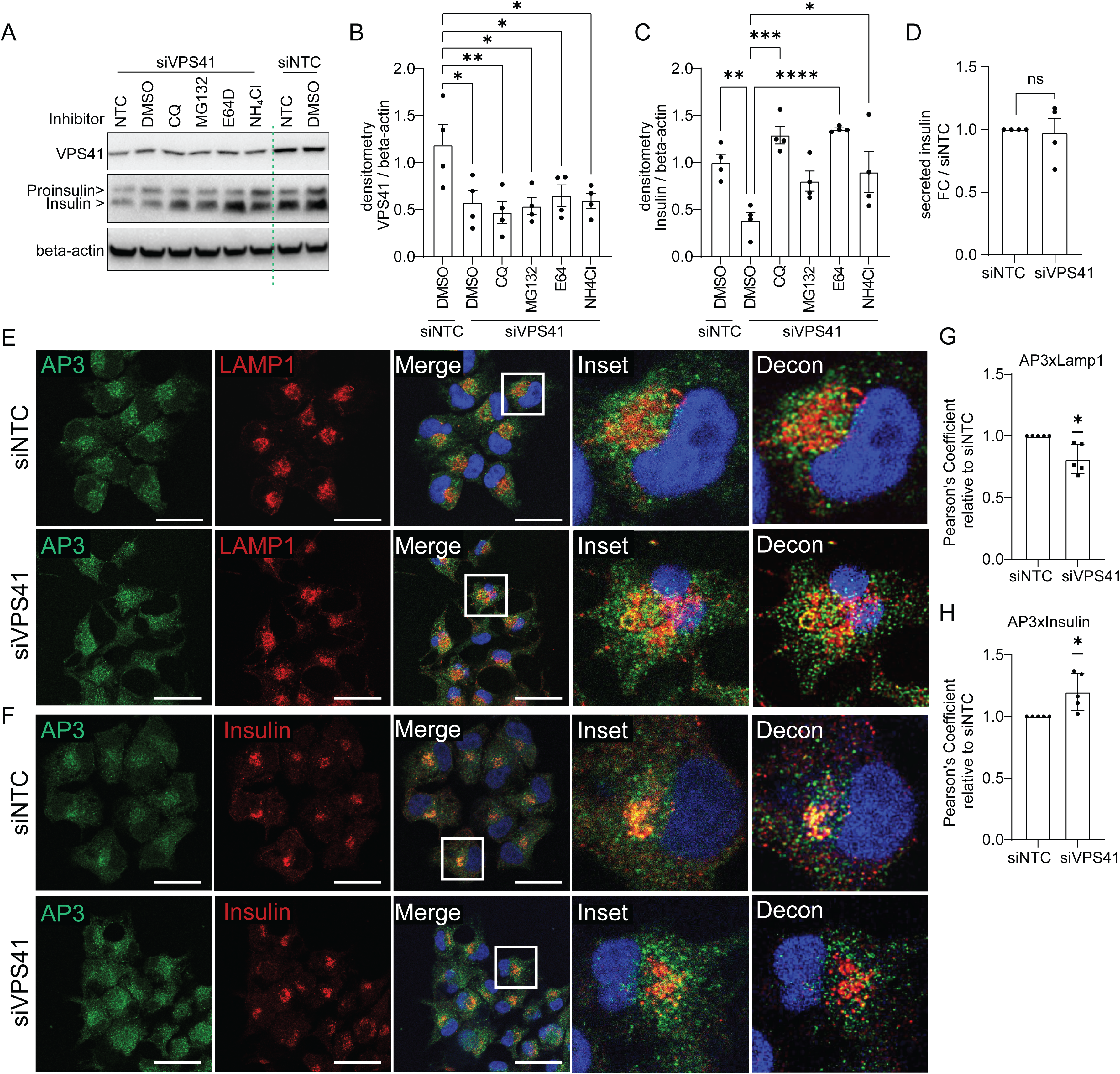
Pretreatment with lysosomal inhibitors rescues insulin in VPS41KD INS1 cells. Representative western blot **(A)** and quantified VPS41 **(B)** and insulin **(C)** expression relative to beta-actin in INS1 cells pre-treated for 18 hours with DMSO, 2µM chloroquine, 100nM MG132, 2µM E64D, or 1mM ammonium chloride (NH_4_Cl), 24 hours post-transfection with VPS41 or NTC siRNA. Insulin measured by ELISA in the media of INS1 cells 24 hours post-transfection with VPS41 or NTC siRNA **(D)**. Representative images of immunofluorescent staining of AP3 and LAMP1 **(E)** and AP3 and insulin **(F)** and corresponding Pearson’s coefficients assessing colocalisation **(G and H)** at 24 hours post-transfection with VPS41 or NTC siRNA. 15 – 20 cells analysed per experiment per condition and each dot represents one experiment. Data expressed as mean ± SD relative to siNTC, *p < 0.05 by one sample t-test..

VPS41 binds to AP3[14], which is essential for insulin granule biogenesis[27]. We hypothesized that the acute loss of VPS41 might be redirecting AP3 from budding insulin granules to lysosomes. To test this, we performed immunofluorescent staining of AP3 and insulin, as well as AP3 and LAMP1, in INS1 cells with VPS41 siRNA-mediated knockdown, 48 hours post-transfection. We observed a significant increase in colocalisation of AP3 with LAMP1, as indicated by Pearson’s coefficient, compared to non-targeting controls (Figure 4E and G). Concurrently, we observed a significant decrease in colocalisation of AP3 and insulin in VPS41 KD compared to non-targeting controls. (Figure 4F and H).

### VPS41 depletion concurrently triggers dedifferentiation of beta-cells through PDX1 downregulation

Previous studies on VPS41 deletion in neuroendocrine cells have reported defective autophagy pathways due to destabilization of the HOPS complex [17], including the nuclear translocation of transcription factor TFE3 and the dissociation of mTORC1 from lysosomes[17]. Our current findings in the INS1 cell line at 24h post-VPS41 knockdown show similar results, with decreased association of mTORC1 and LAMP1 (Figure 5A), and increased nuclear localisation of TFE3 (Figure 5B).

**Figure 5.**
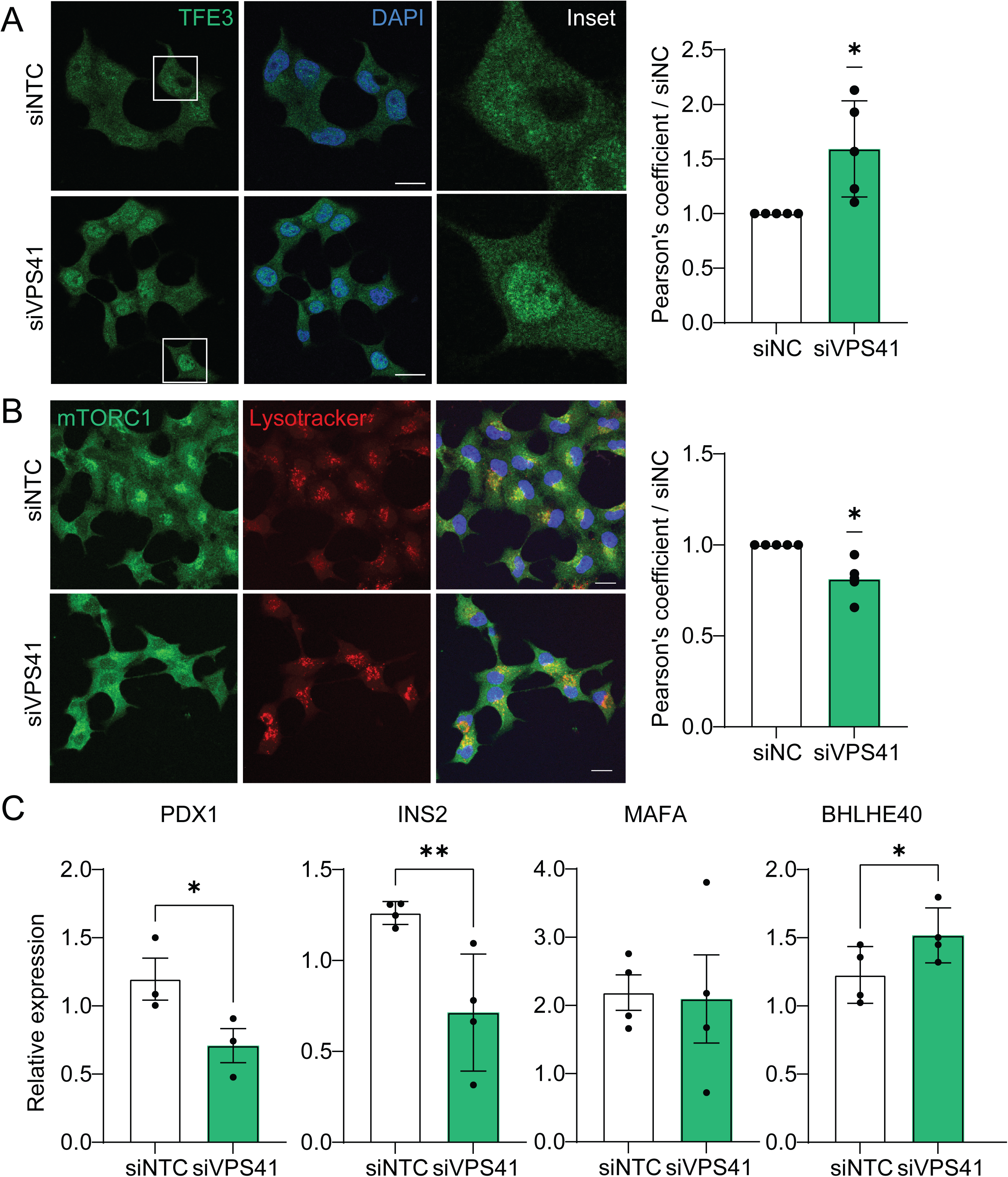
Nuclear localisation of TFE3 associated with downregulation of PDX1 in VPS41KD INS1 cells. Representative immunofluorescent images and quantification of TFE3 and DAPI **(A)** and mTORC1 and lysotracker **(B)** colocalisation in INS1 cells 24 hours post-transfection with VPS41 or NTC siRNA. Relative mRNA gene expression of *Pdx1*, *Ins2*, *Mafa,* and *Bhlhe40* in INS1 cells 24 hours post-transfection with VPS41 or NTC siRNA **(C)**. Data expressed as mean ± SD, *p < 0.05, **p < 0.01 by one sample t-test or unpaired t-test.

In pancreatic beta-cells, TFE3-mediated inhibition of insulin gene expression is particularly notable. Although the precise mechanism is not well-defined, it likely involves activation of the repressor BHLHE40[28], which inhibits MAFA/PDX1 binding to enhancer regions[29]. Consistent with this, we observed significant downregulation of *Pdx1* and *Ins2* mRNA in INS1 cells 24 hours post-VPS41 knockdown, though surprisingly *Mafa* levels remained unchanged, potentially due to differences in timing, as PDX1 regulates MAFA expression[30]. Importantly, we also noted a significant upregulation of the repressor *Bhlhe40* (Figure 5C). Given the downregulation and loss of HOPS complex expression in VPS41 knockdown cells (Figure 3H), these observations suggest a potential link between VPS41-mediated HOPS function, and loss of pancreatic beta-cell identity.

## Discussion

### VPS41 is indispensable to the beta-cell

We first demonstrated the importance of VPS41 in beta-cells through our characterisation of VPS41 INS1 KO cells and beta-cell specific VPS41KO mice[20]. Our studies revealed that genetic deletion of VPS41 in beta-cells led to impaired glucose-stimulated insulin secretion and a dramatic reduction in total insulin content. In mice, this resulted in a distinct diabetic phenotype characterized by insulin insufficiency without associated insulin resistance or immune response. In the current study, we sought to distinguish the effects of chronic insulin deficiency from those of acute VPS41 depletion to gain a deeper understanding of VPS41’s role in both *in vitro* and *in vivo* settings.

VPS41 functions as a coat protein with AP-3 during the early stages of insulin granule budding at the *trans-*Golgi network[14] and is also a core component of the HOPS complex. Notably, there are no functional redundancies for VPS41; while dissolution of HOPS complex in VPS39 KD cells leaves an intact pool of VPS41 that still regulates insulin secretion[20], VPS41 deletion itself is catastrophic for beta-cell function. In our current study using an siRNA knockdown model, we aimed to differentiate the effect of chronic VPS41-induced insulin deficiency from those of acute VPS41 loss-of-function. Despite over 80% VPS41 protein depletion at 72 hours post-siRNA treatment, we observed no defects in glucose-stimulated insulin secretion, although total insulin content was significantly reduced. This indicates a potential disconnect between VPS41’s role in these two processes. It is possible that siRNA-mediated depletion preferentially affects the HOPS-associated pool of VPS41, similar to observations in VPS39 KO cells, though this has not been confirmed. Interestingly, re-expression of a VPS41 variant with a single nucleotide polymorphism (S284P) that corresponds to a clinically relevant human variant (S285P, [17]) in VPS41 KO INS1 cells was able to restore both insulin secretion and insulin content without rescuing HOPS function[20]. This raises intriguing questions about how this HOPS-independent pool of VPS41 contributes to insulin granule biogenesis and exocytosis, which remains an area for further investigation.

### VPS41 loss causes insulin degradation in the beta-cell

Our current study of acute VPS41 depletion reveals that the primary defect is the rapid degradation of insulin. Insulin granule biogenesis relies on the coordinated trafficking of membrane and luminal proteins to aggregate proinsulin and other cargo within immature secretory granules[31]. Both VPS41 and its binding partner AP3 have been implicated in this critical process[14].

Crucially, our study found that pre-treatment with lysosome inhibitors such as chloroquine or the cysteine protease inhibitor E64D was effective in rescuing insulin content in VPS41KD cells. Interestingly, while reductions in total proinsulin and insulin content were observed with chloroquine treatment as measured by ELISA (Supplemental Figure 4G), these changes were not detected by Western blotting. This discrepancy may be attributed to chloroquine-mediated downregulation of proinsulin synthesis[32]. Despite this, the data confirm that the rapid degradation of mature insulin occurs through lysosomal compartments. Therefore, it is likely that either microautophagy or crinophagy – processes involving the direct delivery of insulin granules or their contents to lysosomes – may be erroneously activated in the absence of VPS41.

To our knowledge, the rapid loss of insulin content within hours of depleting a single protein has not been previously observed. Strikingly, even partial depletion of VPS41 leads to significant insulin loss at early as 24 hours post-transfection (Figure 6C). This finding is underscored by the age-dependent loss of insulin in VPS41KO mice (Figure 2E) and is particularly noteworthy in light of a recent study indicating that VPS41 mRNA transcription is downregulated in mouse pancreatic beta-cells under chronic hyperglycaemia[33]. Interestingly, VPS41 KD cells exhibit a phenotype similar to nutrient-starved cells with mTORC1 inhibition coupled with TFE3 nuclear localisation. This suggest that TFE3-driven modulation of the master regulator PDX1 may downregulate insulin transcription[28], potentially through BHLHE40-mediated repression of PDX1. *Pdx1* is crucial for the development of insulin-positive beta-cells and maintenance of beta-cell identity[34]. Consistent proinsulin production is essential for beta-cell identity, as evidenced by proinsulin processing defects in dual *Ins1* and *Ins2* KO mice that cannot be rescued by insulin gene re-expression[35]. Ultimately, we data shows that VPS41 deletion acutely disrupts degradation pathways while depleting insulin stores. Moreover, the absence of VPS41 leads to chronic insulin-deficiency and gradual dedifferentiation of beta-cells, which is reflected in persistent and worsening hyperglycaemia with age. A caveat to our *in vivo* findings is the use of an embryonically-deleted VPS41 beta-cell model, in which chronic changes from a lack of VPS41 during development cannot be extricated. Future studies that may inform VPS41 function include the inducible deletion of VPS41 within beta-cells in adult mice, as well as the inclusion of transcriptomics in beta-cell specific VPS41KO islets, and the measure of VPS41 expression within non-diabetic and diabetic human islets.

**Figure 6.**
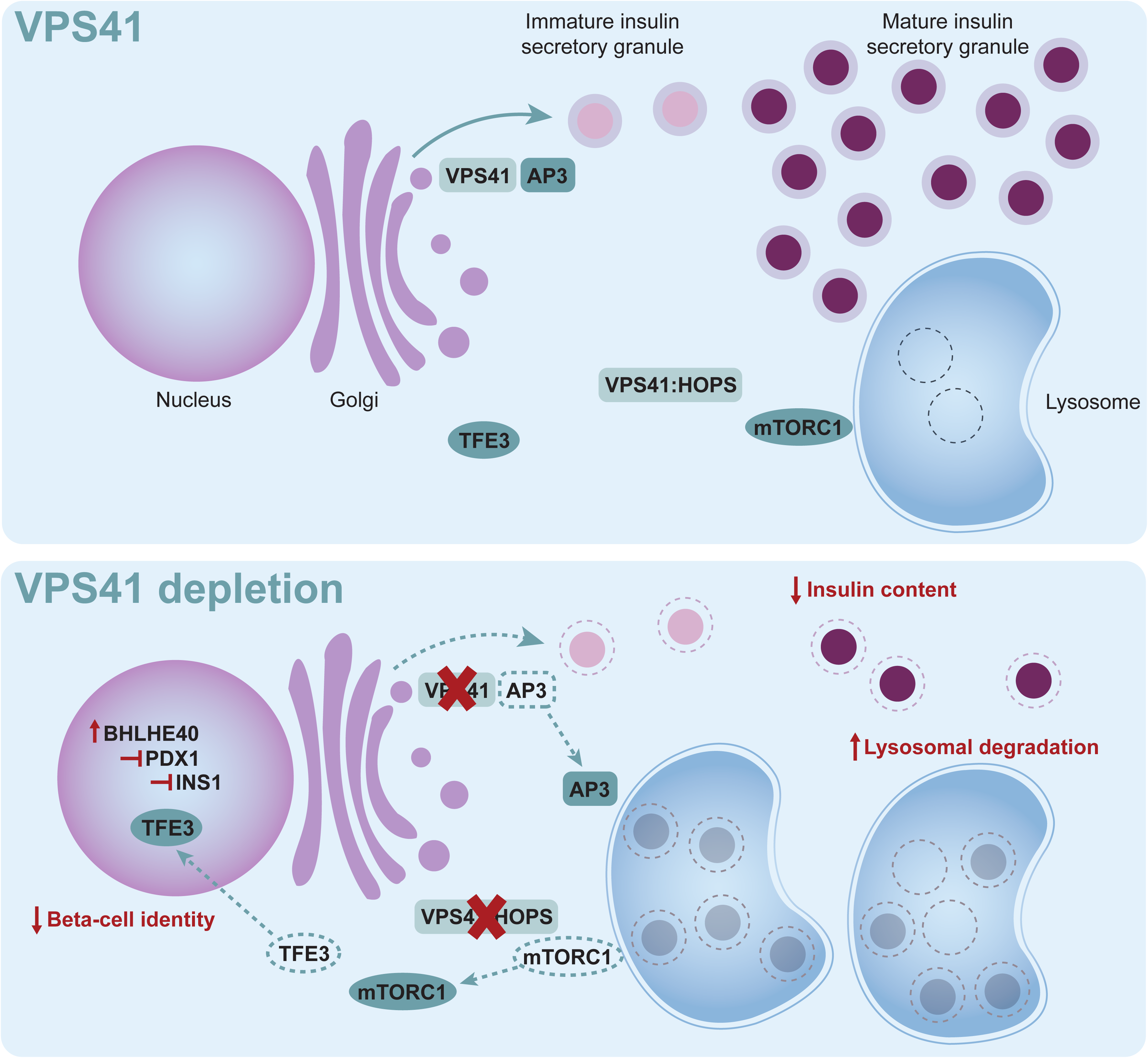
Schematic depicting VPS41-depletion dependent loss of insulin granule stores and beta-cell identity.

These findings highlight the critical role of VPS41 in maintaining insulin granule content, and suggest likely through an AP3-dependent role in insulin granule formation[14]. We also expose a potential HOPS-dependent role in preserving beta-cell identity through TFE3-regulation of *Pdx1* (Figure 6). More broadly, our results emphasize the delicate balance between insulin storage and degradation in the beta-cell and its importance for overall whole body homeostasis.

## Acknowledgements

We would like to thank Elizabeth Lawfull and Jane Calderwood from Australian BioResources (Garvan Institute for Medical Research) for their assistance with mouse ovariectomy procedures, Dr. Ben Crossett (SydneyMS, University of Sydney) with the processing and run of VPS41KD INS1 cell LC-MS/MS samples, and Dr. Ross Laybutt for IRE1-alpha and CHOP antibodies.

## Author Contributions

Conceptualization, MAK, BY, CSA; Methodology, MAK, BY, CSA; Validation, YA, BY; Formal Analysis, BY, YA; Investigation, BY, YA, CB, MG, PS, GV, ML; Resources, MAK, Data Curation, BY, YA; Writing – Original Draft, BY; Writing – Review & Editing, BY, YA, HW, CSA, MAK; Visualization, BY; Supervision, MAK; Project Administration, MAK, BY; Funding Acquisition, MAK. MAK is the guarantor of this work and having access to the data of this study, takes responsibility for the contents of this article.

---

**Supplemental Figure 1.**
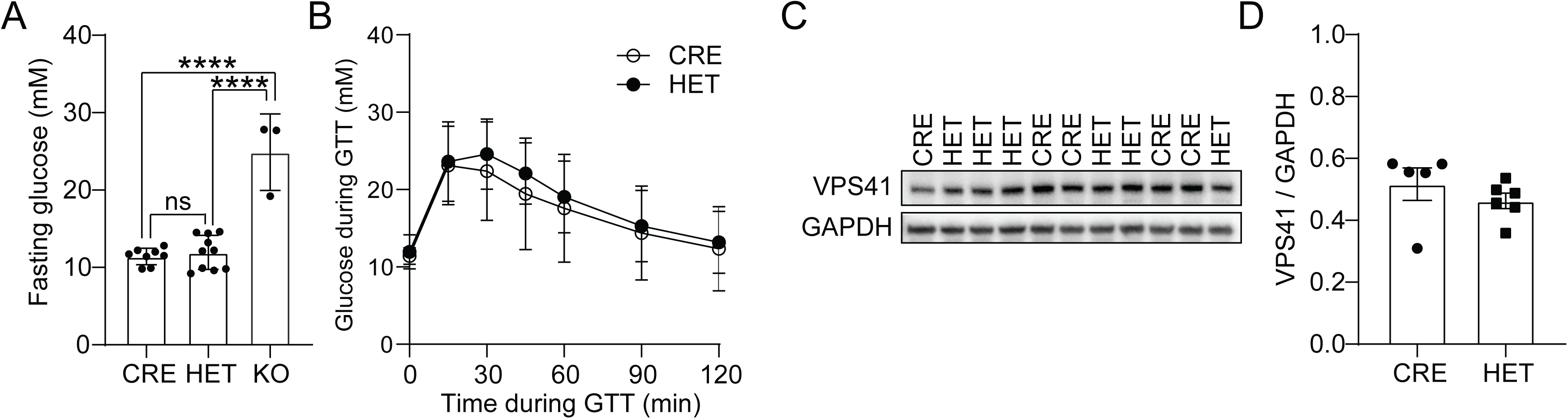

**Supplemental Figure 2.**
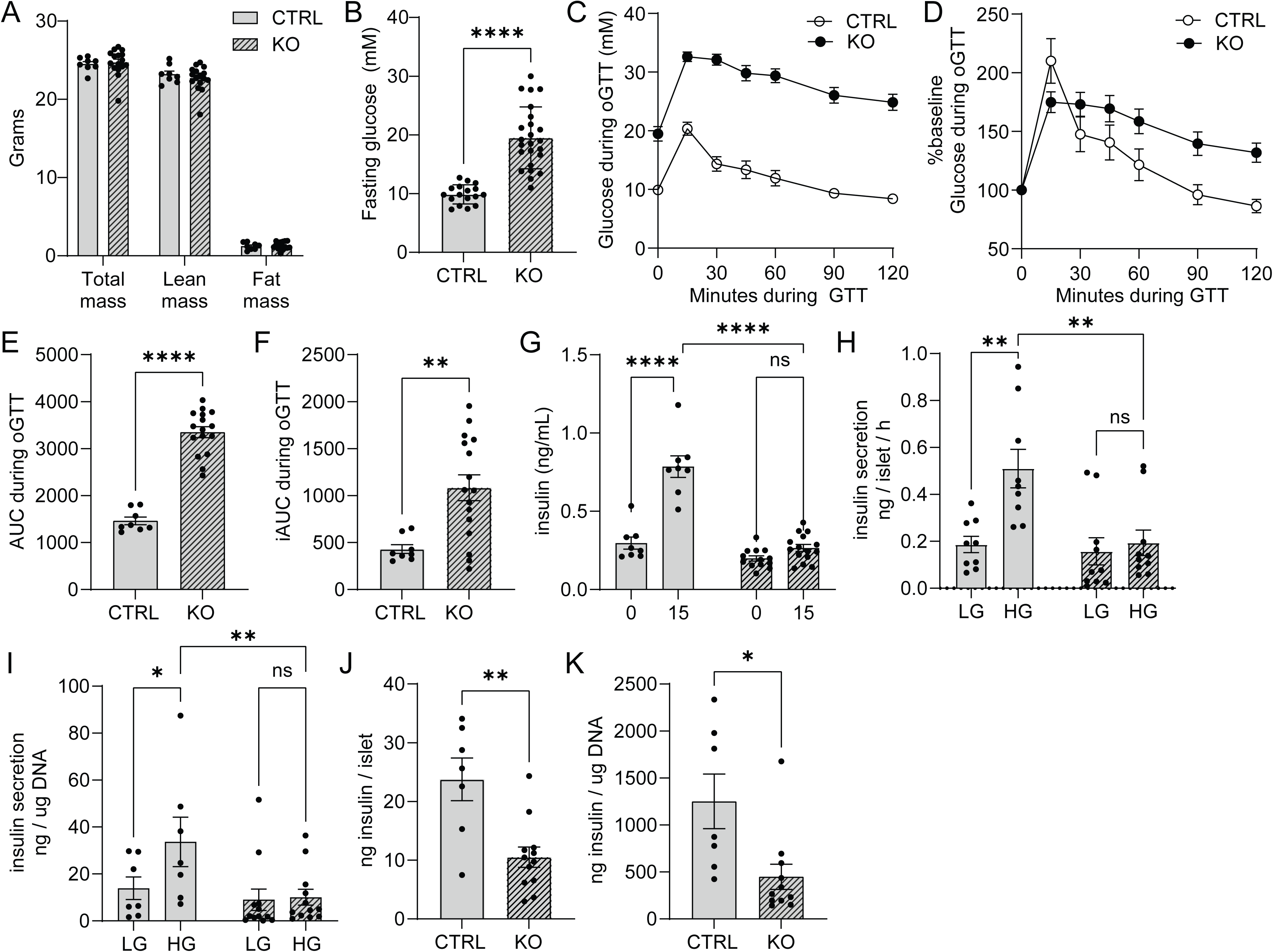

**Supplemental Figure 3.**
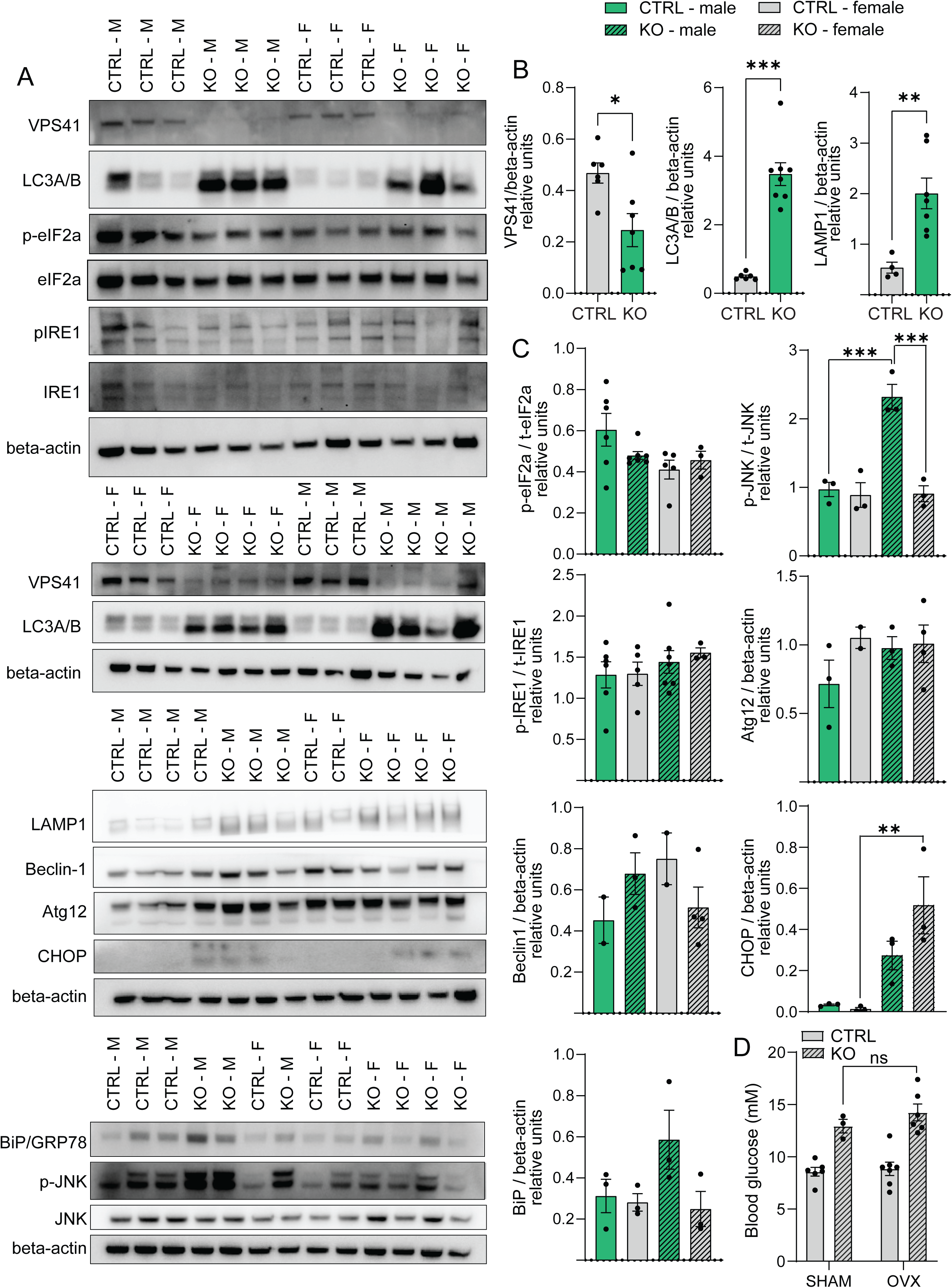

**Supplemental Figure 4.**
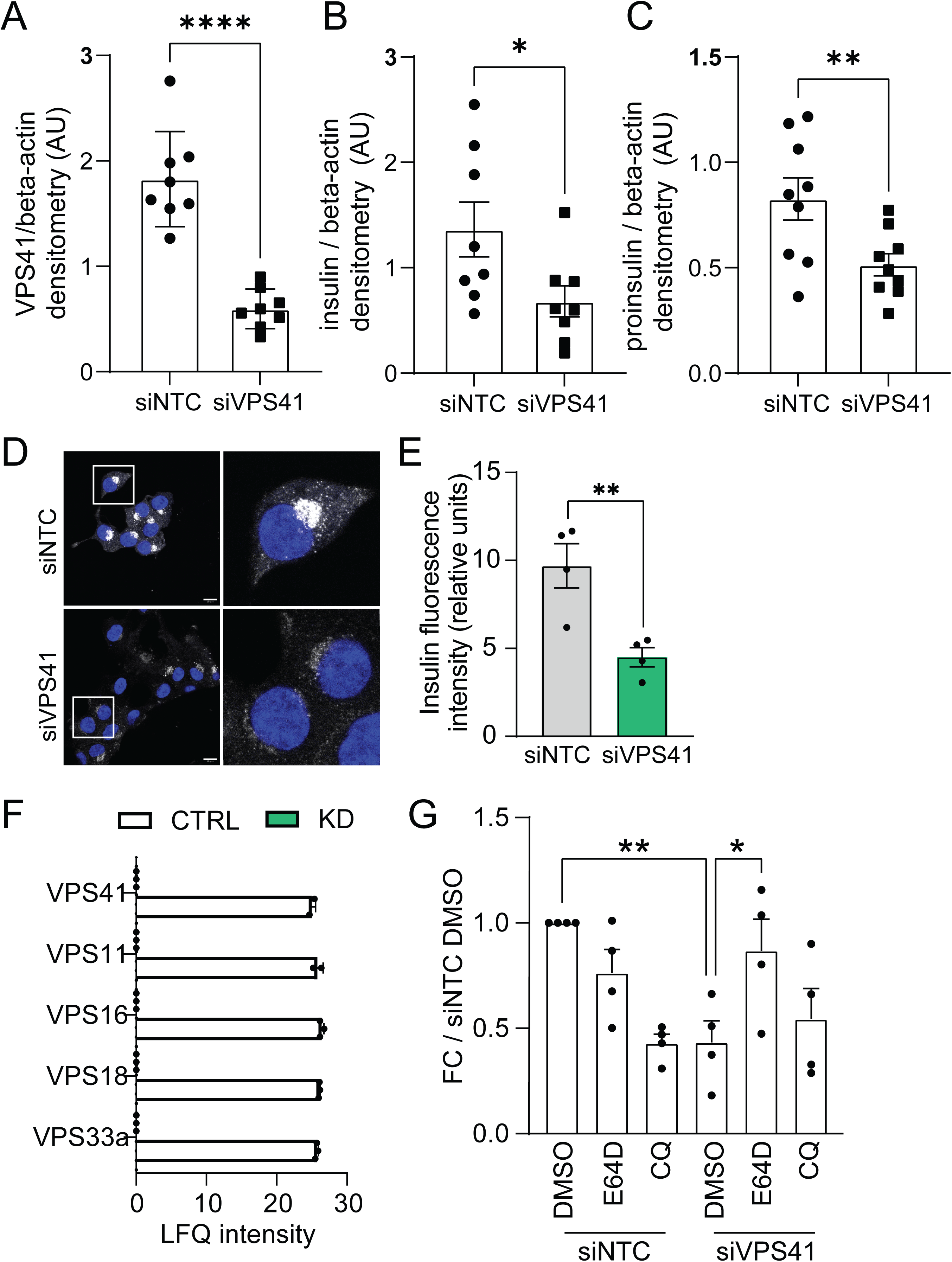

## References

1. Yau, B., et al., Targeting the insulin granule for modulation of insulin exocytosis. Biochem Pharmacol, 2021. 194: p. 114821.

2. Schnell, A.H., I. Swenne, and L.A. Borg, Lysosomes and pancreatic islet function. A quantitative estimation of crinophagy in the mouse pancreatic B-cell. Cell Tissue Res, 1988. 252(1): p. 9–15.

3. Vasiljević, J., et al., The making of insulin in health and disease. Diabetologia, 2020. 63(10): p. 1981–1989.

4. Boland, B.B., C.J. Rhodes, and J.S. Grimsby, The dynamic plasticity of insulin production in β-cells. Mol Metab, 2017. 6(9): p. 958–973.

5. Lee, Y.H., et al., β-cell autophagy: Mechanism and role in β-cell dysfunction. Mol Metab, 2019. 27s(Suppl): p. S92–s103.

6. Pearson, G.L., et al., A Selective Look at Autophagy in Pancreatic β-Cells. Diabetes, 2021. 70(6): p. 1229–1241.

7. Marsh, B.J., et al., Regulated autophagy controls hormone content in secretory-deficient pancreatic endocrine beta-cells. Mol Endocrinol, 2007. 21(9): p. 2255–69.

8. Pasquier, A., et al., Lysosomal degradation of newly formed insulin granules contributes to β cell failure in diabetes. Nat Commun, 2019. 10(1): p. 3312.

9. Goginashvili, A., et al., Insulin granules. Insulin secretory granules control autophagy in pancreatic β cells. Science, 2015. 347(6224): p. 878–82.

10. Shvarev, D., et al., Structure of the HOPS tethering complex, a lysosomal membrane fusion machinery. Elife, 2022. 11.

11. Pols, M.S., et al., hVps41 and VAMP7 function in direct TGN to late endosome transport of lysosomal membrane proteins. Nat Commun, 2013. 4: p. 1361.

12. van der Beek, J., et al., CORVET, CHEVI and HOPS - multisubunit tethers of the endo-lysosomal system in health and disease. J Cell Sci, 2019. 132(10).

13. Lin, X., et al., RILP interacts with HOPS complex via VPS41 subunit to regulate endocytic trafficking. Sci Rep, 2014. 4: p. 7282.

14. Asensio, C.S., et al., Self-assembly of VPS41 promotes sorting required for biogenesis of the regulated secretory pathway. Dev Cell, 2013. 27(4): p. 425–37.

15. Schoppe, J., et al., AP-3 vesicle uncoating occurs after HOPS-dependent vacuole tethering. Embo j, 2020. 39(20): p. e105117.

16. Jiang, D., et al., Arabidopsis HOPS subunit VPS41 carries out plant-specific roles in vacuolar transport and vegetative growth. Plant Physiol, 2022. 189(3): p. 1416–1434.

17. van der Welle, R.E.N., et al., Neurodegenerative VPS41 variants inhibit HOPS function and mTORC1-dependent TFEB/TFE3 regulation. EMBO Mol Med, 2021. 13(5): p. e13258.

18. Sanderson, L.E., et al., Bi-allelic variants in HOPS complex subunit VPS41 cause cerebellar ataxia and abnormal membrane trafficking. Brain, 2021. 144(3): p. 769–780.

19. Steel, D., et al., Loss-of-Function Variants in HOPS Complex Genes VPS16 and VPS41 Cause Early Onset Dystonia Associated with Lysosomal Abnormalities. Ann Neurol, 2020. 88(5): p. 867–877.

20. Burns, C.H., et al., Pancreatic β-Cell-Specific Deletion of VPS41 Causes Diabetes Due to Defects in Insulin Secretion. Diabetes, 2021. 70(2): p. 436–448.

21. Thorens, B., et al., Ins1(Cre) knock-in mice for beta cell-specific gene recombination. Diabetologia, 2015. 58(3): p. 558–65.

22. Rohli, K.E., et al., ER Redox Homeostasis Regulates Proinsulin Trafficking and Insulin Granule Formation in the Pancreatic Islet β-Cell. Function (Oxf), 2022. 3(6): p. zqac051.

23. Harney, D.J., et al., Dietary restriction induces a sexually dimorphic type I interferon response in mice with gene-environment interactions. Cell Rep, 2023. 42(6): p. 112559.

24. Yau, B., et al., Proteomic pathways to metabolic disease and type 2 diabetes in the pancreatic islet. iScience, 2021. 24(10): p. 103099.

25. Yau, B., et al., A fluorescent timer reporter enables sorting of insulin secretory granules by age. J Biol Chem, 2020. 295(27): p. 8901–8911.

26. Okita, N., et al., Modified Western blotting for insulin and other diabetes-associated peptide hormones. Sci Rep, 2017. 7(1): p. 6949.

27. Sirkis, D.W., R.H. Edwards, and C.S. Asensio, Widespread dysregulation of peptide hormone release in mice lacking adaptor protein AP-3. PLoS Genet, 2013. 9(9): p. e1003812.

28. Pasquier, A., et al., TFEB and TFE3 control glucose homeostasis by regulating insulin gene expression. Embo j, 2023. 42(21): p. e113928.

29. Tsuyama, T., et al., Hypoxia causes pancreatic β-cell dysfunction and impairs insulin secretion by activating the transcriptional repressor BHLHE40. EMBO Rep, 2023. 24(8): p. e56227.

30. Sachdeva, M.M., et al., Pdx1 (MODY4) regulates pancreatic beta cell susceptibility to ER stress. Proc Natl Acad Sci U S A, 2009. 106(45): p. 19090–5.

31. Germanos, M., et al., Inside the Insulin Secretory Granule. Metabolites, 2021. 11(8).

32. Chatterjee, A.K. and H. Schatz, Effect of chloroquine on biosynthesis, release and degradation of insulin in isolated islets of rat pancreas. Diabetes Res Clin Pract, 1988. 5(1): p. 9–15.

33. Cheruiyot, A., et al., Sustained hyperglycemia specifically targets translation of mRNAs for insulin secretion. J Clin Invest, 2023. 134(3).

34. Gao, T., et al., Pdx1 maintains β cell identity and function by repressing an α cell program. Cell Metab, 2014. 19(2): p. 259–71.

35. Ramzy, A., et al., Insulin Null β-cells Have a Prohormone Processing Defect That Is Not Reversed by AAV Rescue of Proinsulin Expression. Endocrinology, 2022. 163(6).

